# Differential degradation of RNA species by autophagy related pathways in plants

**DOI:** 10.1101/793950

**Authors:** D. Hickl, F. Drews, C. Girke, D. Zimmer, T. Mühlhaus, J. Hauth, K. Nordström, O. Trentmann, H.E. Neuhaus, T. Fehlmann, A. Keller, M. Simon, T. Möhlmann

## Abstract

An important function of the plant vacuole is the recycling of the delivered proteins and RNA by autophagy. We provide the first plant vacuolar small RNome by isolation of intact vacuoles from Barley and Arabidopsis, subsequent RNA purification and Next Generation Sequencing. In these vacuolar sRNomes, all types of cellular RNAs were found including those of chloroplast origin, suggesting a bulk-type of RNA transfer to, and breakdown in vacuoles. ATG5 is a major representative of autophagy genes and the vacuolar RNA composition in corresponding knockout plants differed clearly from controls as most chloroplast derived RNA species were missing. Moreover, the read length distribution of RNAs found in ATG5 mutants differed to control samples, indicating altered RNA processing. In contrast, vacuolar RNA length and composition of plants lacking the vacuolar RNase2 (*rns2-2*), involved in cellular RNA homeostasis, showed minor alterations, only. Our data therefore suggests that mainly autophagy components are responsible for selective transport and targeting of different RNA species into the vacuole for degradation. In addition, mature miRNAs were detected in all vacuolar preparations, however in ATG5 mutants at much lower frequency, indicating a new biological role for vacuolar miRNAs apart from becoming degraded.

## Introduction

Vacuoles fulfil many functions in plant metabolism. They are responsible for the maintenance of turgor pressure and they are storage compartments for sugars, organic acids, amino acids and ions [1,2]. Furthermore, vacuoles are also a site for breakdown of cellular macromolecules. The role in RNA degradation was assigned to vacuoles based on the observation of 80% of total cellular RNase activity in this compartment [3]. This was further supported by others [4, 5] and later, evidence for the presence of RNA-oligonucleotides in isolated vacuoles from cultured tomato cells was presented [6]. Bariola [7] identified RNS2, a member of the T2 RNase family from Arabidopsis as vacuolar RNase and Hillwig [8] describe this enzyme as necessary for ribosomal RNA decay and found that corresponding knockout mutants showed a constitutive autophagy. In the same year, with ENT1 (equilibrative nucleoside transporter 1) a vacuolar transport protein was identified, responsible for the export of RNA breakdown products in form of nucleosides [9].

Based on these findings it is clear that plant vacuoles are important sites for cellular RNA catabolism. However, the types of RNA delivered to the vacuole and the identity of catabolic enzymes are so far unknown.

Macroautophagy (termed autophagy in the remaining of the manuscript) is a process that delivers unwanted cellular material and organelles to the vacuole for degradation [10]. In yeast, autophagy was identified as mechanism involved in bulk RNA degradation induced by starvation, pointing to the importance of rRNA turnover e.g. as nitrogen source under stress conditions [11]. In line with this, Arabidopsis plants lacking the vacuolar RNase2 (RNS2) show constitutive autophagy, even under normal growth conditions, emphasizing the role of RNA recycling for plant homeostasis [8,12].

Bassham and MacIntosh [12] state in the Abstract of their review on autophagy mediated degradation of ribosomes that “Ribosomes are essential molecular machines that require a large cellular investment, yet the mechanisms of their turnover are not well understood in any eukaryotic organism. Recent advances in Arabidopsis suggest that plants utilize selective mechanisms to transport rRNA or ribosomes to the vacuole, where rRNA is degraded and the breakdown products recycled to maintain cellular homeostasis” [12,p.1].

To address this topic we developed a protocol, allowing quantitative and qualitative analysis of the vacuolar RNome. Therefore, we have isolated vacuoles from Arabidopsis and towards a general validity in addition from Barley (*Hordeum vulgare*) as a monocot species with agronomical significance. In a novel approach we sequenced the RNome of this organelle to obtain information on the type of RNA at single sequence resolution. Furthermore, we included mutants lacking components of the autophagy process (ATG5) and the RNA catabolism (RNS2) in our analysis.

## Results

In our project we aimed for the identification of RNA species inside plant vacuoles at single sequence resolution. We chose Barley (*Hordeum vulgare*) as an agronomically important monocot plant and Arabidopsis as a general plant model. Although, detailed protocols for the isolation of highly pure vacuoles for both species exist, we had to develop a strategy for the investigation of vacuolar RNA composition. Thus, subsequent to vacuole isolation, RNA was extracted, converted into Illumina compatible libraries and subjected to deep sequencing, followed by bioinformatic analysis.

### Barley vacuoles contain short RNA fragments mainly belonging to chloroplast RNAs

For vacuole isolation, primary leaves from 8-9 day old Barley (*Hordeum vulgare* type “Barke”) plants were harvested at the beginning of the light period, following available protocols (see Methods) [13]. Purified vacuoles were microscopically inspected and found to be intact and free of contaminating protoplasts (Figure 1 A). To obtain first indications for the presence of RNA species inside vacuoles, these were incubated with the RNA specific SYTO® RNASelect™ Green Fluorescent cell stain. Using this dye a clear green fluorescence from the vacuolar lumen was observed (Figure 1B-D). Since the presence of ribosomal RNA (rRNA) in plant vacuoles was shown before [8,14], we checked for the presence of rRNA species in our isolated barley vacuoles. For this, RNA extracted from isolated vacuoles was subjected to northern blot and hybridized with 5S rRNA and 18S rRNA probes. In both cases, small fragments (below 120 nt) were labelled. Smallest fragments were app. 30 nt for the 5S probe and 50 nt for the 18S probe (Figure 1 E). To allow for a better and quantitative estimation of the RNA size distribution, total vacuolar RNA was run on an Agilent small RNA Chip (BioAnalyzer 2100, Agilent Technologies, Santa Clara, USA). The majority of RNA corresponded to a fragment size of 25 to 200 nt (Figure 1F). Intact ribosomal RNA was absent from the sample, supporting a high purity and low contamination of the isolated vacuoles. Following our aim for a description of the vacuolar RNome at single sequence level resolution, a RNA library was constructed and subsequently Illumina sequencing was performed on a HiSeq 2500 (Illumina, San Diego, USA), and obtained reads were trimmed removing reads <15 nt and mapped to the available Barley annotated genome [15]. The distribution of the top 12 of all mapped reads to gene loci is shown in the pie chart (Figure 1G). Most prominent were reads mapping to the chloroplast 16S rRNA gene (gene id = HORVU2Hr1G062290) with 52% of top 12 reads. Three loci encode cell wall associated hydrolases (HORVU2Hr1G062280, HORVU7Hr1G064930, HORVU6Hr1G046740; total of 31% of top 12 reads). At positions 4-6 and 8-12 in our ranking we find photosynthesis associated nuclear genes (phang) rbscL, pSBA, pSBD, and Cp43 (Figure 1G, Supplemental Table S1 (htseq count_barley)).

**Fig. 1.**
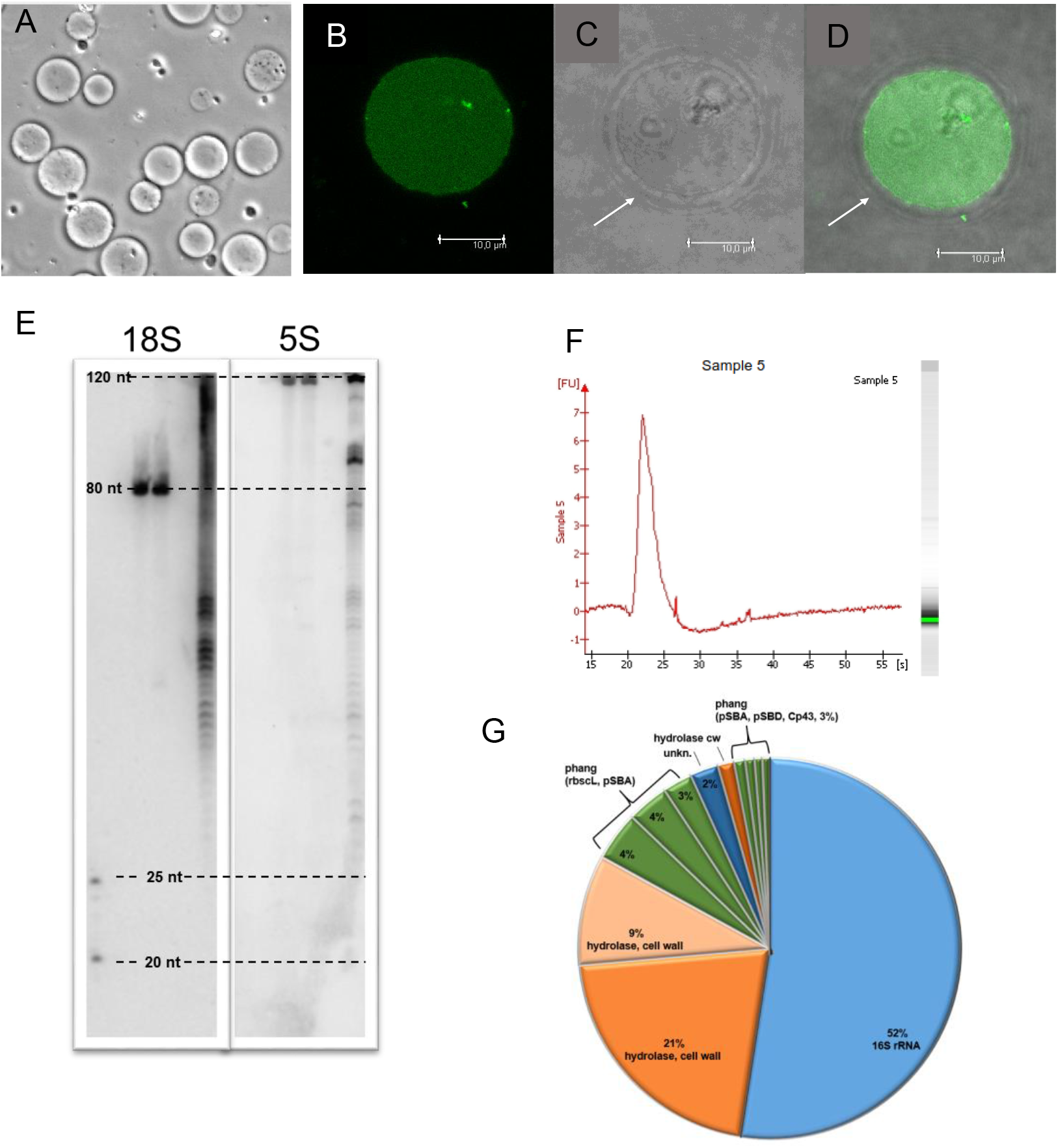
Isolated rRNA from Barley vacuoles with 25 to 200 nt in size is qualitatively classified. (**A**) Vacuoles were isolated from 8-9 day old *Hordeum vulgaris* plants at the beginning of the light period. Microscopical inspection showed pure vacuoles without contaminating protoplasts. (**B-D**) RNA inside the vacuolar lumen was detected by incubation with SYTO® RNASelect™ Green Fluorescent cell stain (B). Brightfield image (C) shows vacuolar intactness and the overlay image (D) reveals RNA fluorescence only from the inside of the vacuole. Tonoplast is indicated by arrowhead (C and D; scale bar = 10 μm). (**E**) Separated RNA was blotted on a nylon membrane and hybridized with 5S rRNA and 18S rRNA probes. **F**) Total RNA isolated from vacuoles was run on a Agilent RNA Pico Chip. The single peak with a width of 20 to 27 seconds at its base represents eluting RNA fragments with a size of 25 to 200 nt. (**G**) Illumina sequencing enabled classification of RNA species. Top 12 identified genes are shown with % frequency of reads. Photosynthesis associated nuclear genes (phang).

### Arabidopsis vacuoles contain short RNA fragments belonging to rRNA, tRNA and mRNA derived from cytosol and chloroplasts

In a second step we aimed to decipher the vacuolar RNome from Arabidopsis. For this we applied two slightly differing protocols for the isolation of vacuoles. A first protocol requires small leaves from plants that have not been singularized after sowing. (referred to as R-type vacuoles) [16]. The alternative method uses rosette leaves from individually grown Arabidopsis plants (referred to as B-type vacuoles) [17].

Based on marker enzyme activities, extravacuolar contaminations were investigated. Uridine diphosphate glucose pyrophosphorylase activity (UGPase, representing cytosol and intact protoplasts) was 3.44%± 0.71 for R-type and 10.8% ± 3.55 for B-type vacuoles, compared to intact protoplasts. Contamination of purified vacuoles by chloroplasts was detected by Glycerine aldehyd-3-phosphate dehydrogenase activity and was below the detection level for R-type vacuoles and 3.54%± 0.78 for B-type vacuoles compared to intact protoplasts (Figure 2A).

**Fig. 2.**
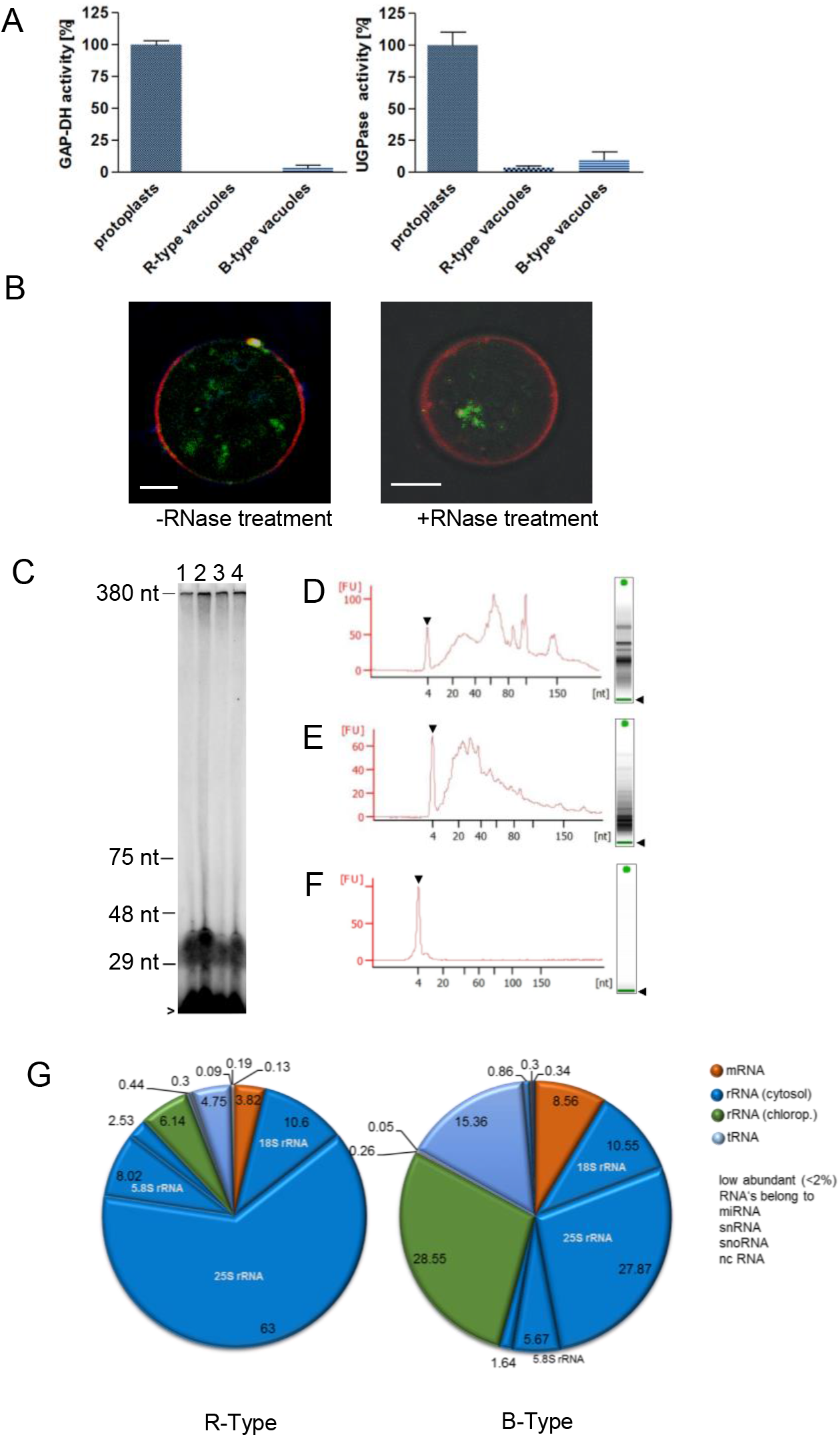
Purity, intactness and RNA composition of R and B-Type vacuoles from Arabidopsis. Plants were grown on soil for 4 to 6 weeks with dense sowed seeds for isolation of R-type vacuoles and 3 to 4 plants per pot for isolation of B-type vacuoles. (**A**) R-type and B-type vacuoles are almost free of extravacuolar contaminations. Marker enzyme activity was used for verification of plastidic (Glycerine aldehyd-3-phosphate dehydrogenase (GAPDH)) or cytosolic contamination (Uridine diphosphate glucose pyrophosphorylase (UGPase)). Enzymatic activity in protoplasts was used as control and set to 100%. (**B**) Tonoplast protects vacuolar RNA against extravacuolar RNase activity. Intravacuolar RNA from B-type vacuoles was stained with the RNA specific dye SYTO® RNASelect™ (green), whereas the tonoplast was visualized by the lipophilic dye FM4-64 (red). Treatment with RNase prior to staining had no influence on RNA signal (right; scale bar = 4 μm). (**C**) RNA from R-type vacuoles was non treated (lane 1), with 10 μg ml^−1^ (lane 2), 0.5 mg ml^−1^ (lane 3) and during cell lysis while preparation with 10 μg ml^−1^ RNase A treated (lane 4). Afterwards RNA was radioactive endlabelled and separated by RNA-PAGE. For size determination 29 nt, 48 nt, 75 nt and 380 nt marker oligos were run on the same gel. Excessive free [γ-^32^P]-ATP is indicated by arrowhead. (**D-F**) B-type vacuoles were non treated (D) or incubated for 1 h with 10 μg ml-1 RNase A, followed by 1 h incubation with 400 μg ml^−1^ Proteinase K (E). Isolated RNA was separated on RNA chip. RNase A activity was verified by 15 min incubation of 10 μg ml^−1^ RNase A with 400 ng protoplastic RNA (F). (**G**) Illumina sequencing and mapping enabled classification of RNA species. RNA composition in R- and B-type vacuoles differs in plastidic rRNA, t-RNA and mRNA content.

To check for the presence of RNA within the vacuoles, the preparations were incubated with the RNA specific dye SYTO® RNASelect™ and the lipophilic dye FM4-64. The staining reveals presence of RNA inside B-type vacuoles (Figure 2B), whereas no fluorescence signal was detected in R-type vacuoles. Treatment of intact B-type vacuoles with RNase A prior to staining did not diminish the SYTO RNASelect signal from the vacuolar interior (Figure 2B) indicating protection of RNA against external RNase by the tonoplast.

For a quantitative readout of probable RNA contaminations R-type vacuoles were treated with different concentrations of RNase A and RNase A plus proteinase K. Latter controls will inactivate residual RNase A activity that could interfere with RNA isolation from vacuoles. Isolated vacuolar RNA was then end-labelled using P^32^-CTP and separated on RNA PAGE. In all lanes most intense labelling was observed at a size of 29 to 48 nt. No obvious difference between treated and untreated samples was observed supporting our previous investigation, that RNA was protected from external RNase by the tonoplast (Figure 2C).

For B-type vacuoles we used RNA chip analysis to detect effects of RNAse A treatment on the size distribution of isolated vacuolar RNA. Whereas a few larger RNAs could be removed by RNase A treatment (Figure 2D), the smaller fraction of RNA was fully protected against degradation (Figure 2E). As a positive control for the functionality of the RNase A treatment, lysed vacuoles were treated with this enzyme and as expected, all RNA was completely degraded (Figure 2F).

### Comparing RNA contents from the two different vacuolar preparations

In a next step we subjected isolated RNA from R-type and B-type vacuoles to small RNA library preparation. RNA was isolated from at least three individual preparations of vacuoles. To obtain a higher read number and thus a better coverage during the mapping procedure, Illumina sequencing was performed on a HiSeq 2500 (Illumina, San Diego, USA), with 6 M and 9 M sequences obtained per library, see methods for details of the procedure. Obtained sequences were trimmed, removing sequences <15 nt and mapped to the Arabidopsis genome (Araport11) with corrections given in Supplemental file S4. Results from one sequencing run for each vacuole type are shown. In R-type vacuoles, the majority of sequencing reads were mapped to cytosolic rRNA loci (84.14%, Figure 2G). With 63% of all reads, the 25S species of cytosolic rRNA is the dominating one. Chloroplast rRNAs, tRNAs and mRNAs follow with 6.14%, 4.75% and 3.82%, respectively (Figure 2G). 0.71% of all reads are distributed among small RNA species as mi-RNA, sno-RNA, nc-RNA and sn-RNA (Figure 2G). Comparing these results with those from B-type vacuoles, the following observations can be made: First, cytosolic rRNA is also the dominant RNA species in B-type vacuoles, with 45.72% of all reads (Figure 2G). The reduction compared to R-type vacuoles is mainly due to reduced reads for 25S rRNA, dropping from 63% to 27.87 % in B-type vacuoles. Second, the largest increase in read number was observed for chloroplast rRNA species, accumulating 28.55% of all reads. Similarly, tRNA (15.36%) and mRNA (8.56%) revealed increased read numbers (Figure 2G). Small RNA types accumulate to 1.55% in sum. For details of read distributions please refer to Table S2.

### AtRNS2-T-DNA insertion lines showed reduced vacuolar adenylate contents and activity

RNS2 was identified as vacuolar RNase involved in a housekeeping function in RNA homeostasis. Knockout mutants of *RNS2* (*rns2-1* and *2-2*) have been described exhibiting RNA accumulation and constitutive autophagy [8,18]. However, the effect of a loss of RNS2 on vacuolar RNA composition and that of breakdown products and intermediates has not been analyzed so far. *Rns2-2* (SALK_069588) was described as real knockout line of *RNS2* but *rns2-1* as expressing a C-terminal truncated version of *RNS2* [19]. Therefore, we characterized another independent T-DNA insertion line, *rns2-3* (SAIL_338_G11). RT-PCR confirmed the production of a null mutant by T-DNA insertion (Figure S1). The level of autophagosomes in *rns2-2, rns2-3* was monitored by staining with monodansylcadaverine (MDC) of seedling roots and counting in the elongation zone as described before [8] and the number of autophagosomes was clearly increased in both lines compared to the wildtype (Figure 3A). Whereas WT roots showed 3.2 ± 0.6 autophagosomes per cell, in *rns2-2* and *rns2-3* the corresponding numbers were 6.8 ± 0.9 and 6.3 ± 0.8 (Figure 3A). This observation is in line with previous reports and corroborates *rns2-3* like the other RNS2 mutants exhibit constitutive autophagy. T2 type RNases typically cleave RNA in an endo-nucleolytic way, via the catabolic intermediates 2’, 3’-cNMP and 3’NMP. When we incubated RNA with vacuolar content, both intermediates, here shown as 2’, 3’-cAMP and 3’AMP could be detected by HPLC. Clearly, vacuolar content from *rns2-2* and *2-3*, normalized to the same activity of the vacuolar marker enzyme alpha mannosidase, produced significantly less breakdown products. Corresponding production of 2’, 3’-cAMP was reduced to 28% for *rns2-2* and *2-3*, and adenosine to 23% and 37% for *rns2-2* and *2-3*, respectively (Figure 3B,C). From this reduced production a RNA hydrolytic activity of wildtype vacuolar content of 17 nmol RNA nucleotides h^−1^ was calculated (Figure 3D). Activity in vacuoles from *rns2-2* plants was reduced to 5 nmol RNA nucleotides h^−1^ and 7 nmol RNA nucleotides h^−1^ in *rns2-3* (Figure 3 D). When we inspected the contents of both breakdown products in freshly isolated vacuoles (R-type), a similar picture emerged. Here, the wildtype contents of 2’,3’-cAMP was 69 ± 3.5 nmol U^−1^ mannosidase and only 13.2 ± 2.2 nmol U^−1^ mannosidase in *rns2-2* and 14.8 ± 5 nmol U^−1^ mannosidase in *rns2-3* (Figure 3 E). No detectable amounts of 3’-AMP or adenine appeared indicating the presence of vacuolar nucleotidase-activity and the absence of vacuolar nucleosidase activity. However, adenosine was found inside vacuoles, accumulating to 25.3 ± 3.7 nmol U^−1^ mannosidase in WT vacuoles and to only 13 ± 2.1 nmol in *rns2-2* and 11.7 ± 2.7 nmol in *rns2-3* vacuoles (Figure 3E).

**Fig. 3.**
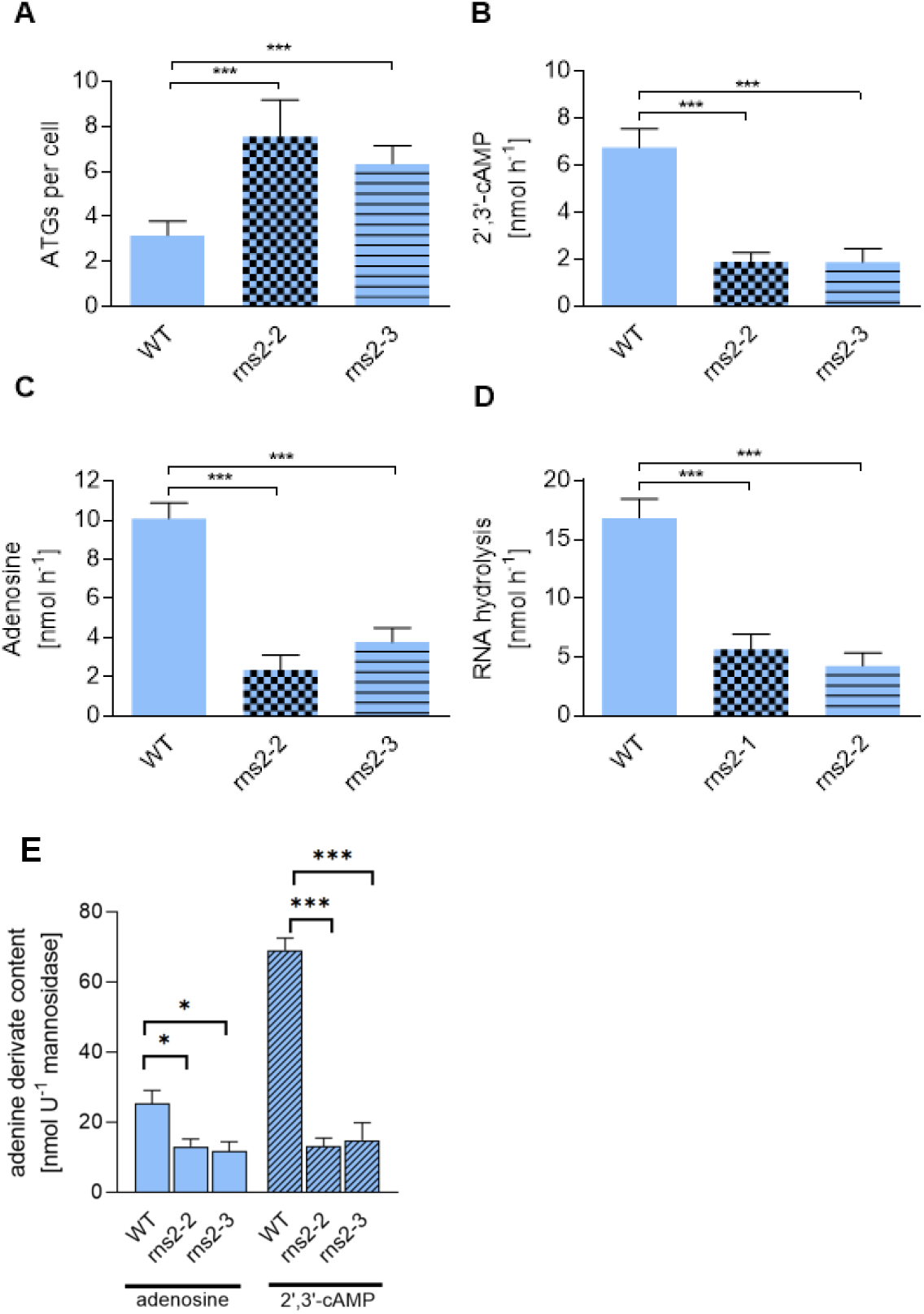
T2 type RNase breakdown products are reduced in RNS2-2 and RNS2-3 mutant plants. (**A**) RNS2-2 and RNS 2-3 exhibit constitutive autophagy. Seedling roots were stained with 0.05 mM monodansylcadaverine (MDC) in PBS for 10 minutes. Observed autophagosomes were counted within elongation zone. (**B-D**) RNA was incubated with vacuolar content from wt and *RNS2* mutants. Produced adenine derivatives 2’,3’-cAMP (B) and adenosine (C) were quantified by HPLC and normalization to vacuolar marker enzyme α mannosidase. (D) By this RNA hydrolysis activity was calculated. (E) Contents of adenosine and 2’,3’ -camp were quantified in different genotypes.

### Comparing the vacuolar RNome from ATG5 and RNS2 mutants

Because of its important role in vacuolar RNA metabolism, a RNS2 mutant (SALK_069588) was compared to wildtype control in respect to the composition of the vacuolar RNome. According to our current understanding, autophagy is the mechanism to deliver cellular macromolecules, damaged organelles and ribosomes to the vacuole for degradation. This involves proteins as well as RNA [12]. ATG5 is a major constituent of the autophagy machinery in Arabidopsis, and thus we included a well characterized ATG5 T-DNA insertion mutant (SAIL_129_B07) in our analysis [20].

ATG5 knockout mutants are characterized by an altered metabolism in prolonged darkness, leading to the inability to survive several days of darkness. When we performed such an experiment, ATG5 knockout plants did not survive a 6 days dark treatment, whereas wildtype and both RNS2 knockout mutants could recover completely (Figure 4A).

**Fig. 4.**
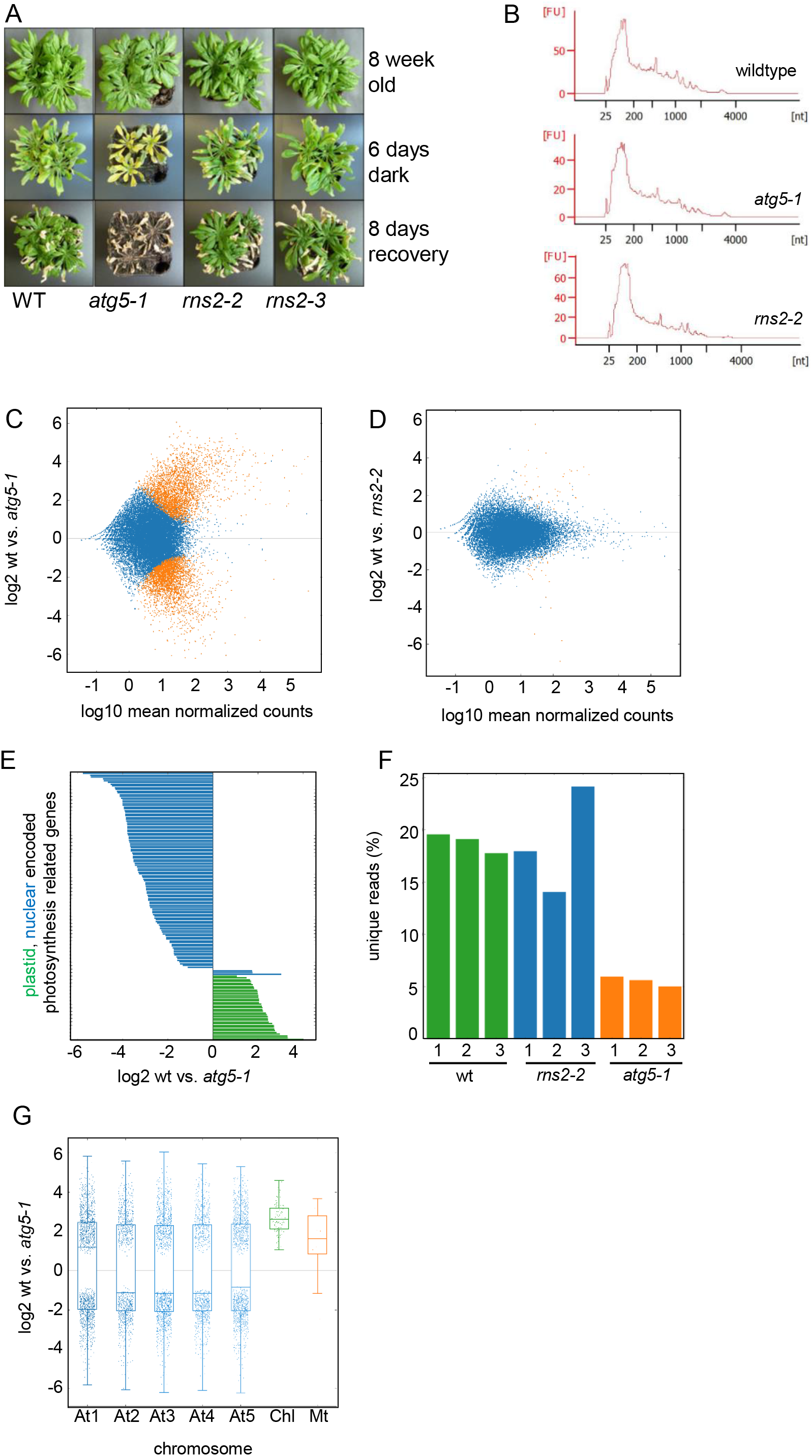
Chloroplast encoded photosynthesis related gene products are strongly decreased in ATG5 vacuoles. (**A**) ATG5 plants are inable to cope with dark induced carbon starvation. Eight week old plants, grown under short day conditions (10 h light, 14 h dark), were transferred to complete darkness for 6 days. Replacing the plants to light recovered wildtype, *rns2-2* and *rns2-3*, but not *atg5-1* plants. (**B**) RNA isolated from B-type vacuoles are 25 to 200 nt in size. No alterations observed in *atg5-1* and *rns2-2*. (**C-D**) Dseq analysis revealed significant differences (p value <0.05) in read numbers of 5080 mapped loci compared between ATG5 and WT (red dots, C), whereas only 65 mapped loci were significantly altered in read numbers compared to RNS2-2 and wildtype (D). (**E**) Photosynthesis related (p.r.) nuclear genes (Phang) were more abundant in ATG5, but p.r. chloroplast encoded genes were less abundant compared to wildtype. (**F**) Percentage of unique reads in wildtype, RNS2-2 and ATG5. (**G**) Chloroplast encoded gene products are markedly decreased in RNA isolated from ATG5 vacuoles.

For the comparison of the vacuolar RNome, three rounds of vacuole (B-Type) isolations from wildtype and the three mutants (*rns2-2, rns2-3, atg5-1*) were performed. Whereas the purity, intactness and presence of RNA fragments inside vacuoles was shown before (Figure 2), we again compared these parameters for all vacuole isolation experiments. Although slight variations in those parameters occurred between biological replicates, overall no significant differences between vacuoles from different genotypes became apparent (Table S3). This also holds true for the extractable amount of RNA (Table S3). To check for the size distribution of RNAs from the three genotypes, Bioanalyzer runs on Pico-RNA chips were performed. The majority of RNAs showed a size of 25 to 200 nt in all three genotypes with only very few larger RNA species. However, no indication for the presence of intact ribosomal RNA (size > 1000nt) was found in any sample analyzed (Figure 4B). In addition, the RNA size profiles of all three genotypes were similar (Figure 4B).

After library preparation and sequencing as described above, between 22 M and 58 M reads were obtained (Table S3). Subsequently, adapter sequences were removed and sequences < 12 nt in size (< 13 nt for miRNAs) excluded from files used for mapping. Mapping was performed by as given in methods on the Araport11 genome assembly corrected by incompletely annotated ribosomal RNA species according to (Table S4). A high number of reads accumulated on the cytosolic 45S rRNA gene and tRNA genes. Comparing accumulated reads of the 45S rRNA gene illustrated a complete coverage of the gene with the exception of ITS1 and 2 regions (Figure S2). We interpret this as indication for a DNA free sample preparation (Figure S2). Furthermore, the read length distribution markedly differed between ATG5 samples and both, wildtype and RNS2 samples (Figure S2).

A differential analysis with DeSeq2 comparing WT and ATG5 found 5080 gene loci with different read numbers out of a total of 29430 loci with mapped reads (Figure 4C). Similar numbers were obtained when comparing ATG5 and RNS2. However, only 65 gene loci differ between WT and RNS2 (Figure 4D). Detailed results from the differential analysis (p value <0.05) comparing the three genotypes can be found in Table S5.

A more detailed sample analysis revealed marked differences in photosynthesis related genes. Whereas, 37 corresponding chloroplast encoded gene products were 1.06 to 4.02 fold (log2) more abundant in WT vacuoles, 55 from 58 photosythesis associated nuclear genes (phang) were more abundant (1.11 to 5.77 fold (log2) in *atg5-1*; Figure 4E). Among the 65 gene loci differing between WT and RNS2 no phang or chloroplast encoded photosynthesis-related gene loci are present i.e. this observation was unique for the ATG5 samples (Table S5). Instead, mRNAs encoded by genes of functional categories “RNA metabolism” and “Protein degradation” were highly represented (Table S5). When inspecting unique reads, it appeared that *atg5-1* mutants accounted for only 5% of unique reads whereas *rns2-2* and wt exhibited 15-20% unique reads (Figure 4F). This large proportion of missing reads could be attributed mainly to chloroplast genome encoded reads (Figure 4G).

### Vacuoles contain intact miRNAs

Micro RNAs (miRNAs) represent a class of small RNAs exhibiting regulatory functions in plant cells. As we identified these molecules in our vacuolar preparations, we investigated these in more detail and extracted miRNA reads from mapping populations. The relative abundance of miRNAs in our libraries is low, usually below 2% of total reads which is drastically lower as one would expect from cellular sRNA libraries which usually contain 50-80% miRNA reads, especially as we used 5’-monophosphate specific ligation which enriches for canonical miRNAs.

The median of read length of these miRNAs was 21nt for eight out of nine samples, only in case of *atg5-1*, 2 it was 20 nt, thus these reads match the size of mature and functional miRNAs (Figure 5A). For most samples, 75% miRNA mapping reads were between 18 and 21 nt long indicating the presence of mature mRNAs instead of degradation products. These would be apparent in Fig. 5 A as our length trimming of reads still maintains reads above 15 nt. Only *atg5-1*, 1 and 2 exhibited a slightly larger zone (Figure 5A).

**Fig. 5.**
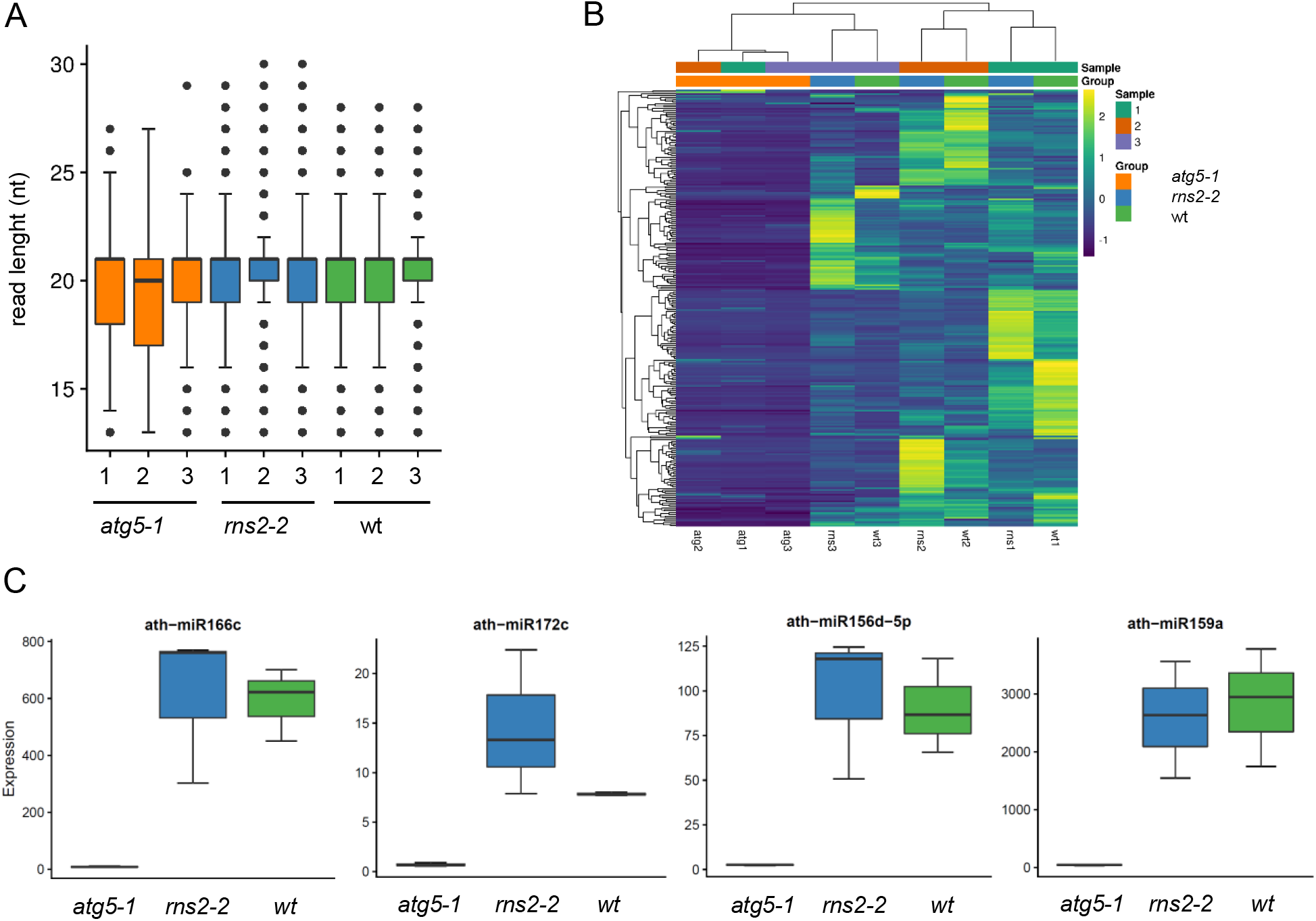
Analysis of miRNAs found in vacuoles from the three different genotypes. (**A**) Size distributon of miRNAs. A median length of 21nt was observed for most samples. (**B**) Cluster analysis of miRNAs indicated lower amounts in ATG5 plants. (C) typical examples of miRNAs with different frequencies in ATG5, RNS2 and WT plants.

A clustering analysis over all 427 miRNAs revealed reduced frequencies for all three ATG5 samples. This is illustrated by selected individual miRNAs miR166c, miR172c, miR156d-5P and miR159a (Figure5C). Although numerous miRNAs have been discovered in animals and plants, many of the corresponding functions in plants are only predicted [21]. Among them is miR166c, which is predicted to target mRNAs coding for HD-Zip transcription factors and thereby regulating organ development and meristem maintenance [22]. Another example is given by miR172c thought to target mRNAs of APETALA2-like transcription factors, thus influencing flowering and floral organ identity [23]. With miR156d, which is predicted to target mRNAs coding for Squamosa-promoter binding protein (SBP) box, we found a further representative of miRNAs modulating floral development [24]. MiR159a, predicted to target MYB transcription factors involved in defense and developmental mechanisms. Herein gibberellic acid and auxin mediated regulation in phosphate starvation and growth take place [22,25,26].

## Discussion

The presence of RNA in vacuoles and corresponding RNase activity is known since long [3,6,7,8]. More recently, RNA delivery to this compartment by autophagy received attention [18,27,12]. In vacuoles from barley seedlings and Arabidopsis rosette leaves, isolated according to established protocols, evidence for the presence of (r)RNA was obtained by us using different techniques: 1. Staining of RNA by Syto-RNA Select in intact vacuoles (Figure 1,2) 2. RNase treatment of isolated vacuoles clearly indicated that RNAs are located inside vacuoles as they were protected against enzymatic degradation. 3. Northern hybridization of RNA purified from intact vacuoles with rRNA specific probes. By this, small sized signals in the range of 20 to 120 nt were detected for vacuolar RNAs from Barley and Arabidopsis (Figure 1, 2).

Cloning of isolated vacuolar RNAs and Illumina sequencing allowed a genome wide identification of RNA reads and a quantification of these. When categorizing the mapped reads, rRNAs were found being most prominent, making up to 92% of all reads found from vacuoles isolated according to Robert [16]. Besides cytosolic rRNAs also chloroplastic (cp) rRNAs were found in differing amounts. In Barley, a similar picture emerged. 52% of the top 12 reads (accounting for 71% of all reads) mapped to the chloroplast 16S rRNA. Furthermore, 7 photosynthesis associated nuclear genes (phang) were identified (12%). This is similar to our results for Arabidopsis. The identification of cell wall associated hydrolases might be more specific for cereal species. Comparing two vacuole isolation protocols, we found that R-Type vacuoles contained only little cp rRNA (6%), whereas B-type vacuoles contained 29% cp rRNA. Barley and R-Type vacuoles were isolated from developing leaves, in contrast, B-type vacuoles were obtained from fully developed, mature leaves. It can be assumed that mature leaves exhibit higher rRNA turnover and contain more aged or damaged chloroplasts prone to degradation [28].

Autophagy is a process involved in the degradation of cellular macromolecules and even organelles [12,14,18,29]. This includes Rubisco-containing bodies (RCBs), ATI1-PS (ATG8-interacting Protein 1) bodies, and small starch-like granule (SSLG) bodies [27,30]. In addition, autophagy independent degradation of damaged chloroplasts has been reported [28,31]. Both pathways, autophagy dependent and independent may contribute to the observed accumulation of RNAs in vacuoles. Besides rRNA other types like t-RNAs, mRNAs and small RNAs (miRNA, snRNA, snoRNA, ncRNA) were found in our analysis. Overall, this clearly points to plants operating a bulk transport of RNA species to the vacuole.

RNS2 was identified as vacuolar RNase of Arabidopsis and corresponding knockout lines were shown to exhibit a longer rRNA half life and constitutive autophagy [8]. In our analysis of two independent RNS2 T-DNA insertion lines we could confirm the presence of an increased autophagosome number in corresponding root cells. At the same time, RNA degradation products were decreased in these lines, as well as the extractable vacuolar RNase activity. However, about 30% of residual RNase activity was found in both RNS2 knockout lines, in line with similar observations by others [19]. This provides evidence for the presence of further RNases in Arabidopsis vacuoles. It was expected to find a higher percentage of non degraded RNAs in RNS2 knockout mutants [12]. However, when we analysed the size of isolated RNAs no difference to wildtype controls or ATG5 mutants became obvious. In addition to our comparison of vacuolar RNA contents between RNS2 and wildtype plants, only 65 genes were identified with significantly altered mapped reads. However, almost no alteration in the read length profile of mapped reads at highly covered loci was observed. We explain the finding of an overall similar RNA size, read profile and RNA composition by a compensatory function of further, so far unknown vacuolar RNases.

In contrast to the minor impact of RNS2 on vacuolar RNA composition, ATG5 showed a huge effect. ATG5 is a main component of the Arabidopsis autophagy system conjugating to ATG12, a ubiquitin like protein tag. The ATG5-ATG12 conjugate is required for autophagosome formation, closure and transfer to the vacuole [32,33]. In line with this, fewer autophagosomes were identified by MDC labelling in *atg5-1* [34] and these mutants are characterized by a massive carbon starvation phenotype and early senescence. When grown on soil, they cannot survive a 5-6 day dark-induced carbon starvation (Figure 4) [32]. Under the same conditions, the effect on wildtype as well as on RNS2 mutants was small. When the RNA composition of *atg5-1* was compared to wildtype, a large amount of genes with altered read number were found. A detailed analysis revealed that most of these reads mapped to the chloroplast chromosome (Figure 4 D). This might be explained by a high selectivity of ATG5 related autophagy for chloroplast RNA. However, as *atg5-1* does not accumulate autophagosomes, the question about the fate of chloroplast RNA in this mutant remains. At least four pathways for the degradation of Chloroplasts or parts thereof exist: 1. Chlorophagy, 2. Piecemeal autophagy leading to formation of rubisco containing bodies (RCB), 3. Chloroplast vesiculation (CV) and 4. The formation of senescence associated vacuoles (SAV). Chlorophagy functions in analogy to mitophagy, describing the selective elimination of dysfunctional mitochondria in yeast and mammals [35]. Similarly chlorophagy can remove entire damaged chloroplasts. Due to photodamage, these chloroplasts were found in wildtype vacuoles, but accumulated in the cytoplasm of ATG mutants [36]. The RCB pathway mobilized stromal proteins to the vacuole [37] and both pathways are autophagy dependent [36]. It is therefore unlikely they play roles in *atg5-1* plants. In contrast, CV represents an autophagy independent way to deliver chloroplast contents to the vacuole in form of small vesicles [31]. A second autophagy independent pathway is defined by SAVFormation. SAV’s represent an independent lytic compartment that does not require fusion with the central lytic vacuole [38,39]. If SAV is favoured in Atg5 plants, we are blind for RNA, which is incorporated in SAVs. Indeed chloroplasts were lost faster in *atg5-1* plants and it was thus supposed this was based on the activation of alternative, autophagy independent breakdown routes [32]. CV and SAV might be good candidates for such alternative routes. However, to the best of our knowledge for none of the before discussed chloroplast degradation pathways RNA has been identified as a target. Therefore, it is hard to predict whether chloroplast RNA breakdown involves CV or SAV or so far unknown, RNA specific pathways.

Next to the smaller fragments of longer RNAs which are likely degradation products, we also detected miRNA reads. Surprisingly, the read length of vacuolar miRNAs is similar to that of mature plant miRNAs, indicating that these are nor degraded but rather being intact molecules.

In general, miRNAs play important regulatory roles in plant developmental processes and the response to nutrient starvation by acting on transcription factors (40). In mammals autophagy itself is target of regulation by miRNAs. [41]. Such regulation has so far not been identified for plants. The median read length of vacuolar miRNAs was 21 nt in eight out of nine samples, matching exactly the typical length of mature, functional miRNAs. Supposedly, miRNAs are prone to degradation in vacuole by autophagy [21]. This might explain the reduced frequency of miRNAs found in ATG5 plants. However, the question appears why intact, mature miRNAs dominate in vacuoles. It is known that miRNAs circulating in the blood stream of mammals are surprisingly stable, possibly by a permanent protection against RNAse by Argonautes (AGO). In plants, the presence of AGO in vacuoles is not known, and we cannot explain right now the biological significance of vacuolar miRNAs. They may represent a reservoir for regulatory miRNAs and could be involved in regulation of plant circulating sRNAs. However, the vacuole itself appears to hold more RNA secretes than just serving as a compartment for their degradation.

## Materials and Methods

### Plants used for analysis and growth conditions

We used wildtype Columbia Arabidopsis plants and T-DNA-insertions lines for *RNS2* (At2g39780) and *ATG5* (At5g17290). These were SALK_069588 (*rns2-2*) SAIL_338_G11 (*rns2-3*) and SAIL_129_b07 (*atg5-1*) [42, 43]. Barley (*Hordeum vulgare* type “Barke”) seeds were a gift from Breun Saatzucht (Herzogenaurach, Germany). *Arabidopsis thaliana* (L.) Heynh. plants (ecotype Columbia) were grown in standardized ED73 (Einheitserde und Humuswerke Patzer, Buchenberg, Germany) soil grown at 120 μmol quanta m^−2^ s^−1^ in a 10 h light/14 h dark regime (short day), temperature 22°C, humidity 60%). Prior to germination, seeds were incubated for 24 h in the dark at 4°C for imbibition. Barley was grown in the same soil under the same growth conditions. RNS2-3 T-DNA insertion mutant analysis as given in Figure S1 was performed with primers listed in Table S6.

### Vacuole isolation

The isolation of *H. vulgare* vacuoles was performed similar to the protocol from Rentsch and Martinoia [13] with modifications in [44,45]. For the isolation primary leaves from 8-9 day old plants were harvested at the beginning of the light period. Isolated vacuoles were either used directly for experiments or frozen at −20 °C for later RNA isolation. The isolation of *A. thaliana* vacuoles (B-type) was performed according to the protocol of Burla [17]. For this we used leaves of Arabidopsis plants singularized and grown under short day conditions for six weeks. The isolation of *A. thaliana* vacuoles (R-type) was performed according to the protocol of Robert [16]. Arabidopsis seeds were sown in pots and grown without singularizaton for 4 to 6 weeks under short day conditions. Plants were harvested before the onset of light.

### RNA preparation

RNA was prepared from isolated vacuoles using chloroform-phenol extraction (peqGOLD TriFast™ FL, Peqlab, Erlangen, Germany) according to the manufacturers advice. The reagent was directly added to frozen vacuoles to inhibit any RNase activity. To enhance RNA yields, glycogen (0.05 − 1 μg μl^−1^) was added to the aqueous phase during subsequent precipitation with EtOH.

### Labelling of nucleic acids

Probes for Northern hybridization were labelled with [α-^32^P]-dCTP using Rediprime II (Amersham, Bucinghapshire, UK) according to the manufacturers instruction. Endlabeling of RNA was achieved with T4-polynukleotide kinase (New England Biolabs) For this, 10 μM RNA was incubated with 1 μl [γ-^32^P]-ATP (150 μCi μl^−1^), 1.5 μl T4-Polynukleotid-Kinase (10 U μl^−1^) in reaction buffer for 1 h at 37 °C. Subsequently RNA was cleaned with QIAquick Nucleotide Removal-Kit (Qiagen, Hilden, Germany). RNA was separated on denaturing gels using formaldehyde (Northern blot) [46] or on high resolution urea 17.5% polyacrylamide gels (PAGE, endlabeled RNA). Quantification was achieved with a MP Storage Phosphoscreen on a Cyclone Phospho-Imager (Perkin-Elmer).

### Enzyme activity and metabolite determination

Alpha mannosidase was quantified according to Pertl-Obermayer [47]. Glycerinaldehyde-3-phosphate-dehydrogenase was determined as given in [48], UDP-glucose-pyrophosphorylase according to Zrenner [49]. Chromato-graphical analysis of adenylates was performed as given in Daumann [50].

### Quantification of autophagosomes

Autophagosomes were quantified as given in Floyd [14]. For each plant line a minimum of 70 cells were inspected. Microscopic analysis was done with a Leica SP5II confocal laser scanning microscope (Leica microsystems, Mannheim, Germany). The following settings were used: (488-nm excitation and 50- to 560-nm detection of emission for SYTO RNA Select, 514-nm excitation and 714- to 754-nm detection of emission for FM4-64 and 405-nm excitation and 474- to 532nm emission wavelength for MDC (monodansyl cadaverine) through a HCX PL APO 63×1.2W water immersion objective).

### Library preparation and Illumina sequencing

RNA libraries were prepared from 70ng vacuole RNA without further size exclusion. Using the „NEBNext® Multiplex Small RNA Library Prep Set for Illumina”-Kit (New England Biolabs, Ipswich, MA, USA) we used diluted adapters and 18 hours for 3’-adapter ligation at 16°C. After 15 PCR cycles, the library was purified from a 6% native TBE-PAGE. Library size and quantity was additionally determined using the Qubit Fluorometer and the Agilent Bioanalyzer DNA HS Chips. Illumina Sequencing was carried out on a HiSeq2500 platform (Libraries WT_1-3, ATG5-1_1-3, RNS2-2_1-3, referred to in Figure 4 and R-type Arabidopsis, B-type Arabidopsis, referred to in Figure 2G) using the RAPID mode for 50 cycles or the Illumina MiSeq (Barley library, referred to in Figure 1G) also for 50 cycles. Reads were demultiplexed and trimmed for adapters using Trim Galore which uses Cutadapt. We eliminated all reads shorter than 13 nt, unless indicated otherwise

### Data analysis

For comparison of R-type and B-type vacuolar RNA contents (Pie charts in Fig. 2G) individual reads were trimmed eliminating reads shorter than 15 nt and aligned using HISAT2 with adapted parameters (−k 20; -secondary). Aligned reads were counted using multicov (bedtools). The percentage of counts for each RNA type was calculated based on the gene annotation.

Barley reads obtained from Illumna MySeq sequencing samples were trimmed eliminating reads shorter than 15 nt and aligned using Bowtie for Illumina from galaxy tools [51]. Counting was performed with htseq counts.

To analyse genotype specific differences in vacuolar RNA composition, reads were processed as follows. First, adapter trimming was performed using TrimGalore (v. 0.4.2, https://github.com/FelixKrueger/TrimGalore). Reads shorter than 13 nt were discarded. Subsequently, reads were mapped against the reference genome using RNA STAR with a maximum intron size of 3000 nt (v. 2.5.2b-0, 52) and quantified using htseq-count (v. 0.6.1, --m intersection-nonempty, 53). Normalization and differential expression analysis among genotypes were carried out using DESeq2 (v. 2.11.40.6, 54). Gene enrichment analysis was calculated conducting a hypergeometric test (55, doi: 10.1093/bioinformatics/btl633) using BioFSharp (v.0.1.0, https://github.com/CSBiology/BioFSharp). All Plots were generated using FSharp.Plotly (v. 1.1.1, https://github.com/muehlhaus/FSharp.Plotly).

For comparison of miRNAs the following procedure was applied. Reads were mapped with up to one mismatch against the reference genome using bowtie 1.2.2 (-a -v1 --best–strata). miRNAs were quantified using featureCounts 1.5.2 and the annotations of Araport 11, while requiring an overlap of at least 80% of the read with the annotation. Fractional counts were assigned to reads mapping to multiple locations (-M --fractional).

Counts were normalized to reads per million (RPM). Clustering of miRNAs was performed with hierarchical clustering using Euclidean distance and complete linkage, and performed on row-scaled RPM normalized counts, while considering only miRNAs that were expressed with at least 5 reads in one sample. The analysis was performed with R 3.5.1 and the plots were generated with pheatmap 1.0.12 and ggplot2 3.1.0.

## Author Contributions

Conceptualization: T.Mö., M.S., C.G., T.Mü, J.H.; Methodology, M.S., C.G., D.H. Investigation: D.H., F.D., C.G. (vacuole isolation, RNA isolation and library preparation) K.N. (Sequencing and Data handling) D.Z., J.H., T.Mü T.F., A. K. (Bioinfomatic and statistic Data analysis) T.Mö., C.G., D.H. wrote the manuscript.

## Acknowledgements

The work was supported by the Forschungsinitiative Rheinland Pfalz (BioComp to T.M.) K.N. was supported by the NBI grant (031L0101D) from the Federal Ministry of Education and Research.

## Supplemental information

Table S1: Read distribution *Hordeum vulgare*

Table S2: Read distribution of Arabidopsis R-type and B-type vacuoles

Table S3: Comparison of “workflow parameters” *wildtype*, *rns2* and *atg5* RNome analysis

Table S4: Araport11 Arabidopsis genome change log

Table S5: DeSeq results comparing *wildtype*, *rns2* and *atg5* RNnome

Table S6: Primers used

Figure S1: *rns2-3* T-DNA insertion mutant analysis

Figure S2: read profiles from 45S rDNA region

