## Supplemental tables and figures for "Differential degradation of RNA species by autophagy related pathways in plants"

Supplemental information

**Table S3** Characterization of isolated vacuoles. Three replicetas of isolated vacuoles from each genotype were prepared and parameters were determined

| **Isolated vacuoles** | WT-1 | WT-2 | WT-3 | ATG5-1 | ATG5-2 | ATG5-3 | RNS2-1 | RNS2-2 | RNS2-3 |
|---|---|---|---|---|---|---|---|---|---|
| α-mann. U $ml^{-1}$ ($10^{-3}$) | 1.42 | 2.06 | 1.66 | 2.43 | 2.24 | 1.20 | 2.47 | 2.48 | 2.50 |
| GAP-DH U $ml^{-1}$ ($10^{-3}$) | 20.82 | 39.7 | 42.0 | 21.6 | 27.2 | 30.5 | 24.6 | 42.3 | 40.8 |
| RNA (µg/ml) | 39.0 | 40.9 | 57.0 | 38 | 59.7 | 44 | 52.2 | 81.5 | 66.1 |
| Protein (µg/ml) | 613.0 | 675 | 667 | 686 | 676 | 643 | 676 | 705 | 746 |
| **Sequencing & data analysis** | | | | | | | | | |
| Input reads (Million, after trimming) | 58 | 42 | 8 | 22 | 31 | 12 | 34 | 33 | 14 |
| Average length (bases) | 26 | 25 | 26 | 26 | 24 | 25 | 26 | 26 | 27 |
| Unique mapped reads (Mio) | 11 | 8 | 1.4 | 1.3 | 1.7 | 0.6 | 6 | 4.6 | 3.4 |
| Mapped to multiple loci (Mio) | 39 | 28 | 5.5 | 19.6 | 27.3 | 10.6 | 23 | 22 | 9.3 |

**Table S6**: Primers used in this study

| *Gene* | s-sequence [5´→3´] | as-sequence [5´→3´] |
|---|---|---|
| knockout screening | | |
| LBb1.3 | ATTTTGCCGATTTCGGAAC | CTGCTTGCAGTGATTCCTTTC |
| LB_3 | TAGCATCTGAATTTCATAACCAATCTCGATACAC | GAGGCCTGTATATTTCCCCAC |
| Ef1α_fwd | GAGACCACCAAGTACTACTGCAC | TGGGTTTTCTAACGCATTGAC |
| Ef1α_rev | GTTGGTCCCTTGTACCAGTCAAG | TCAATGCGTTAGAAAACCCAC |
| RNS2_fwd | ATGGCGTCACGTTTATGTCTTCTCC | CCACCAAGAGAACTGTAAGCG |
| RNS2_rev | TCAAAGAGCTTCTCTTTCTGTTGG | TCTTCAGCAAGACACATGTCG |
| RNS1_RT_fwd | AATCTTGCCTTCTGTCTTCTCTGC | CAAGACCACTCACCACTCCTC |
| RNS1_RT_rev | GTCACAGTATGATCCTGGCCATTG | AGGGTGACATGGATCAGACAG |
| RNS2_RT_fwd | CAAGCTGGCTATGTTGCTTCC | GTCGCAAAAAGCTCGTAGTTG |
| RNS2_RT_rev | GCGTGTATTCCGGCAAACTTACG | TGGTTTTAGCAGCCAAATACG |
| LP/RP 118507 | AGCGAATGGTGGTGCTATATG | TGAGAATCCTTCAGTTCACGG |
| LP/RP 127028C | CAGCAACGGAGACAGAGAGTC | GGGAGGCACTATGAGGTTAGG |
| LP/RP 020074C | CCAAAAGCTCAGGTTCTTGTG | TTTTTATTTTGCTTGTTGGCC |

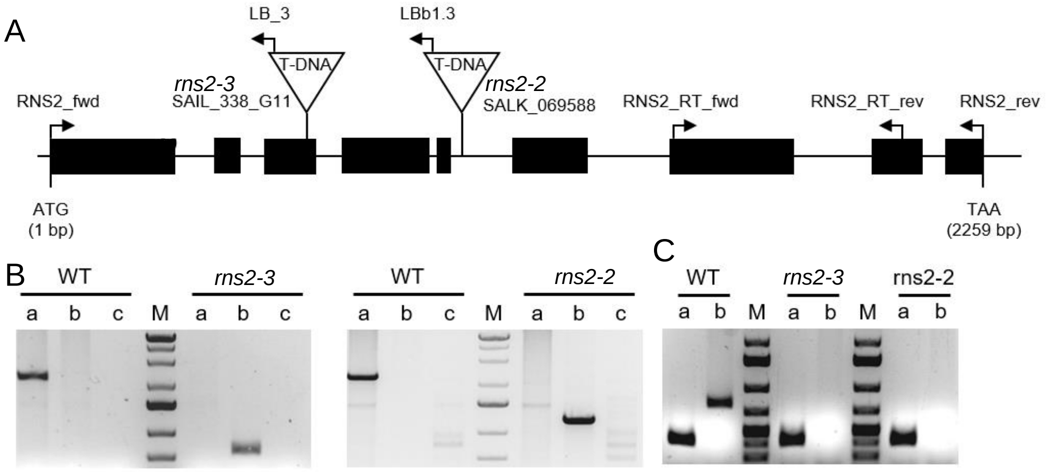


**Fig. S1**. **Identification of RNS 2-2 and RNS 2-3 T-DNA insertion lines. (A)** Schematic view of the *RNS2* gene with position of T-DNA insertion for the lines SALK_069588 (RNS 2-2) and SAIL_338_G11 (RNS2-3), verified by sequencing of specific PCR products. Exon positions are represented by black rectangles, intron positions and untranslated regions by black lines. Primer binding positions are indicated by arrows. (B) Verification of T-DNA insertion by gDNA based PCR analysis. Primer combinations were chosen for gene specific (a) and T-DNA specific amplification products (b and c) in left and middle panel. (C) Complete loss of *RNS2* gene product and quality of cDNA was checked by gene specific primer combination for *RNS2* (b) and the housekeeping gene EF1α in RT-PCR.

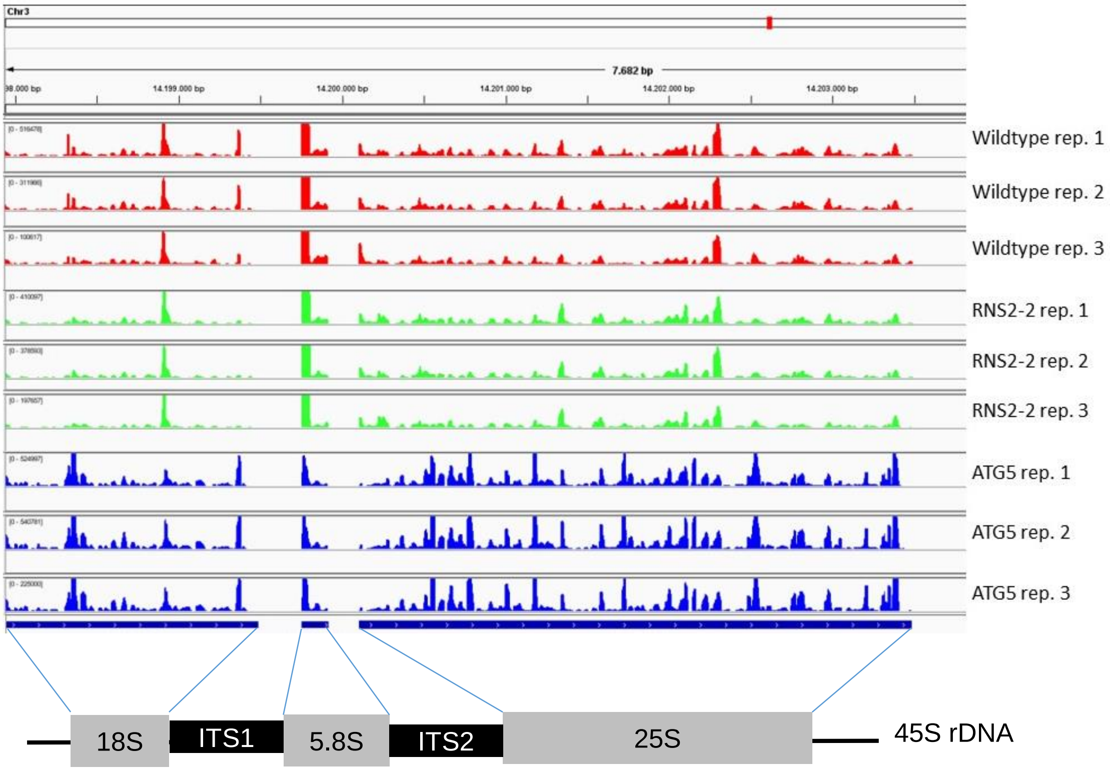


**Fig. S2. Accumulated reads on 45S rDNA in wildtype, *rns2-2* and *atg5-1* plants.** Illustrated by IGV browser Version 2.4.16 downloaded from the Integrative Genomics Viewer homepage, for 3 biological replicates.
